# A Computational Pipeline for Quantifying Kinetochore Morphological Changes in Live Cells

**DOI:** 10.64898/2026.05.26.727517

**Authors:** Jinghui Tao, Vanna Tran, Caleb Rux, Sophie Dumont

**Affiliations:** Bioengineering & Therapeutic Science Department, University of California, San Francisco; Tetrad Graduate Program, University of California, San Francisco; Bioengineering Graduate Program, University of California, San Francisco

## Abstract

To segregate chromosomes kinetochores must resist yet deform under spindle forces. Measuring changes in kinetochore morphology can provide insight into kinetochore structure and function. This remains challenging in live cells because kinetochores are diffraction-limited with irregular, changing shapes. Here, we present a computational pipeline for quantifying kinetochore morphology in live cells, using mammalian cells with fluorescently tagged kinetochore proteins. First, the pipeline tracks, pairs and rotates kinetochores to align with their load-bearing axis. Second, it segments kinetochore signal from background, removing frames with overlapping neighboring kinetochore signals. Third, it provides metrics to define complex, non-Gaussian shape changes: (i) a non-parametric size metric that is more robust than the commonly used full-width-at-half-maximum (FWHM); (ii) analysis to classify common morphological patterns such as asymmetry, low intensity “tails” and multimodality; (iii) a 2D protein1-to-protein2 kinetochore vector as a reporter of structural rearrangements, if two kinetochore proteins were imaged. Finally, we validate the method using simulations, convolving ground-truth objects with the measured point spread function. Although kinetochore shape diversity makes assigning kinetochore size challenging, we show that our metrics better capture kinetochore size and shape changes than FWHM. Together, this pipeline provides a framework for analyzing complex kinetochore shape changes, with potential applications to other small and dynamic cellular structures.

## Introduction

Kinetochores are macromolecular protein complexes built on chromosomes that attach to spindle microtubules to segregate chromosomes at cell division. Kinetochores spend much of their mitotic life under force, and their structure must both resist force and deform under force. In mammals, kinetochores bind the many microtubules that constitute kinetochore-fibers and are made up of over 100 different protein species in multiple copy numbers (Cheeseman & Desai, 2008; Johnston et al., 2010). While kinetochores must maintain their overall structure and their attachment function under force, they must also change shape to facilitate different microtubule attachment states (Magidson et al., 2015) and promote signaling associated with shape (Magidson et al., 2015; Maresca & Salmon, 2009; Uchida et al., 2009, 2021). While we know that mammalian kinetochore shape changes, for example under different attachment scenarios (Cimini et al., 2001; DeLuca et al., 2006), precise, quantitative descriptions of how kinetochore shape and intensity distributions dynamically remodel under different attachment geometries or forces remain limited. Indeed, measuring shape changes remains challenging for small, diffraction-limited objects, especially those, such as the kinetochore, which are deep inside cells, move rapidly, and whose shapes can be complex and that can rapidly change shape over time (Cimini et al., 2001; DeLuca et al., 2006; Kixmoeller et al., 2025; Sacristan et al., 2024). Defining the time evolution of kinetochore morphologies is further challenging in that photobleaching, an issue when creating long timelapse movies, limits the signal to noise ratio of each kinetochore image.

The last two decades have provided great strides in defining the relative position and movement of different protein layers within the mammalian kinetochore (Rago et al., 2015; Wan et al., 2009), including in live cells (Dumont et al., 2012). Imaging of inner and outer kinetochore proteins has shown that kinetochore architecture changes over mitosis (Magidson et al. 2015), and as a function of attachment and force (Dumont et al., 2012; Maresca & Salmon, 2009; Uchida et al., 2009; Wan et al., 2009). However, the shape changes of the inner and outer kinetochore layers themselves, namely the shape changes of the kinetochore intensity distribution, remain understudied. Indeed, given that mammalian kinetochores are small, multilayered structures with inner-to-outer dimensions on the order of ∼100 nm (Brinkley & Stubblefield, 1966; McEwen et al., 1993; Wan et al., 2009), extracting their morphology from live-cell fluorescence images is challenging given the diffraction limit of light. Other quantitative descriptors of kinetochore shape include the full-width-at-half-maximum (FWHM) of the intensity profile (Magidson et al., 2015; Zhou et al., 2025) or using tools to identify ends of kinetochores for length analysis (Cojoc et al., 2016). These measures are appropriate when kinetochore signal intensity over space is approximately Gaussian and unimodal, but can overlook more complex non-Gaussian shape changes that can arise. Both classic and more recent work has reported kinetochore shapes that are complex, irregular and multimodal (Cimini et al., 2001; DeLuca et al., 2006; Kixmoeller et al., 2025; Sacristan et al., 2024). Defining these shapes precisely and quantitatively is a necessary step to understanding how kinetochore structure responds to different attachments and forces, and defining underlying kinetochore structure and organization more broadly.

Here, we present a computational pipeline for quantifying kinetochore morphologies over time in live cell images of mitosis. As a first application and demonstration, we used this pipeline in related work (Tran et al., 2025) to define metaphase kinetochore shape changes, probing how different attachment geometries, force and centromere composition impact inner and outer kinetochore shapes. We now present the pipeline as a broadly usable approach, describing what it does, its mathematical foundation, limits and validation through simulation and controls. In initial image processing steps, the pipeline tracks kinetochores, allows pairing of sisters, aligns kinetochore signals to the force axis, performs signal-to-noise segmentation, filters out potentially overlapping neighbor kinetochores, and projects the kinetochore signal along the force axis to extract 1D intensity profiles. From there, the pipeline delivers metrics defining kinetochores with complex, non-Gaussian shape changes: (i) a non-parametric size metric to define the 2D size (length, width) of irregularly shaped kinetochores; (ii) analysis to classify kinetochore shape asymmetry, low intensity tails, and multimodal intensity peaks; (iii) a 2D vector from kinetochore protein1 to protein2 to test models for what internal structural changes lead to given external shape changes. We validate our approach by simulating kinetochore-like objects of controlled shape, convolving ground-truth masks with the measured point spread function. We show that our size measurement method scales linearly with FWHM for near-Gaussian intensity profiles but remains sensitive to non-Gaussian distributions where the FWHM method overlooks more subtle deformations. Altogether, this pipeline provides a robust and adaptable framework for quantifying kinetochore morphology in live-cell experiments and may be extended to other small, deformable and irregularly shaped cellular structures.

## Results

We now present the computational pipeline we developed for quantifying complex kinetochore morphologies in live cells. We begin by providing an overview of the input data the pipeline can accept, the computational steps in the pipeline and their purpose, and what output metrics the pipeline ultimately delivers (Figure 1). We then describe the image processing steps the pipeline goes through to extract and prepare kinetochore intensity signals for morphological analysis (Figure 2). Then, we the present the metrics we developed to define complex, non-Gaussian shape changes, and how the pipeline extracts them (Figure 3). Finally, we validate our pipeline and define its limits by simulating kinetochore-like objects of controlled shapes and convolving these ground-truth masks with the measured point spread function (Figure 4).

**Figure 1:**
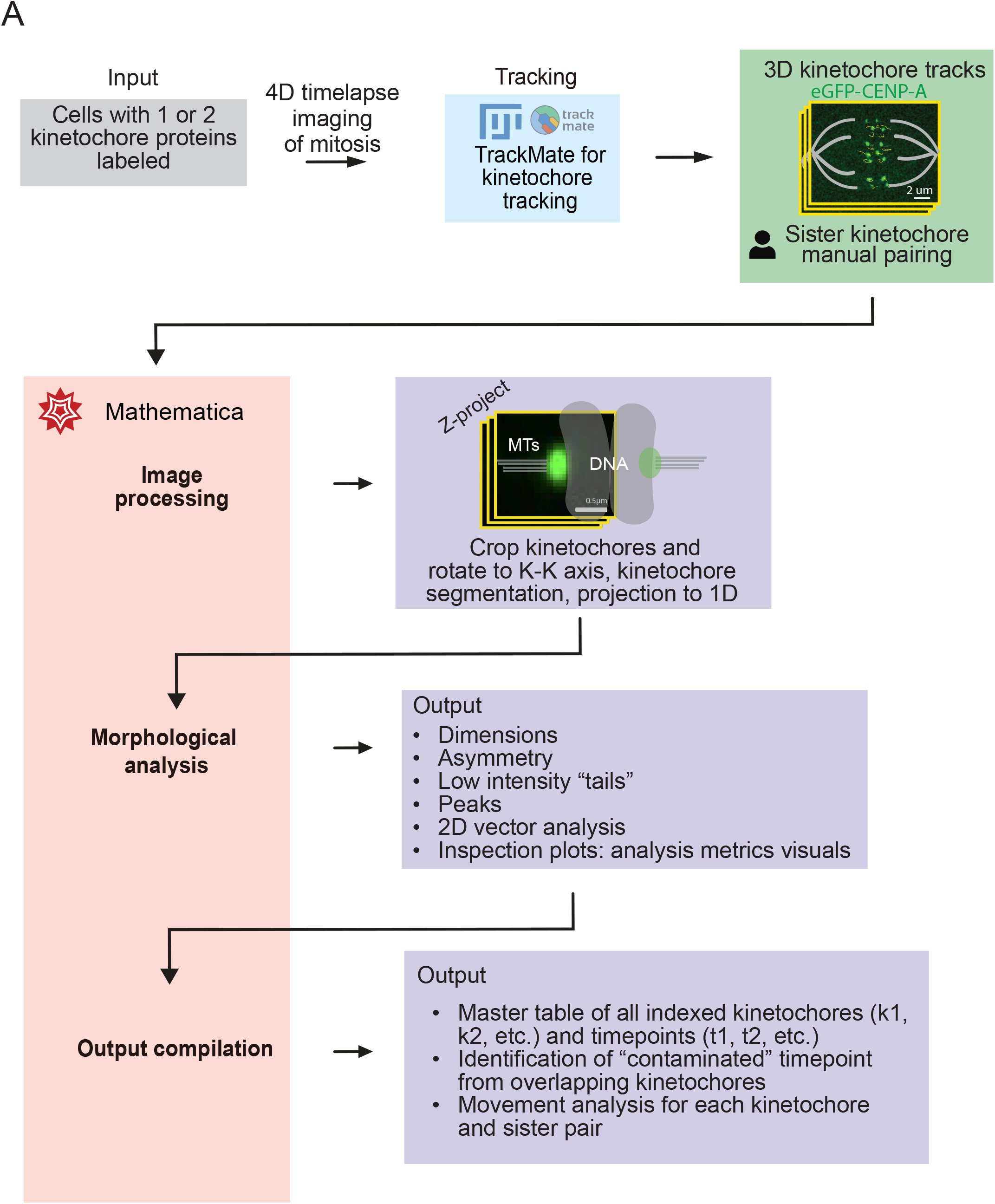
Overview of the computational pipeline for defining complex kinetochore morphologies from live cell movies. (A) Schematic of the workflow for kinetochore morphology analysis. Input (grey box) for the computational pipeline is a timelapse (4D) movie of live cells undergoing mitosis with fluorescently labelled kinetochores. As a first step (blue box), kinetochores are tracked using TrackMate (FiJi) to generate as output (green box) 3D positional tracks over time for each kinetochore which we then manually pair. All further steps are in Mathematica (red box) and with outputs at each step (purple boxes). In Mathematica, each kinetochore image stack is cropped and rotated such that the sister kinetochore-kinetochore axis aligns with the force axis, segmented from the background, and projected to a 1D linescan, (1^st^ purple box). Next, kinetochore intensity signal is analyzed to extract morphological features, including kinetochore dimensions, asymmetry features, low intensity “tails”, peak features, 2D vector between two labelled proteins, and also inspection plots for analysis metric visuals (2^nd^ purple box). The final step is output compilation to compile results of indexed kinetochores and timepoints, identify overlapping kinetochore timepoints, and perform movement analysis (3^rd^ purple box).

**Figure 2:**
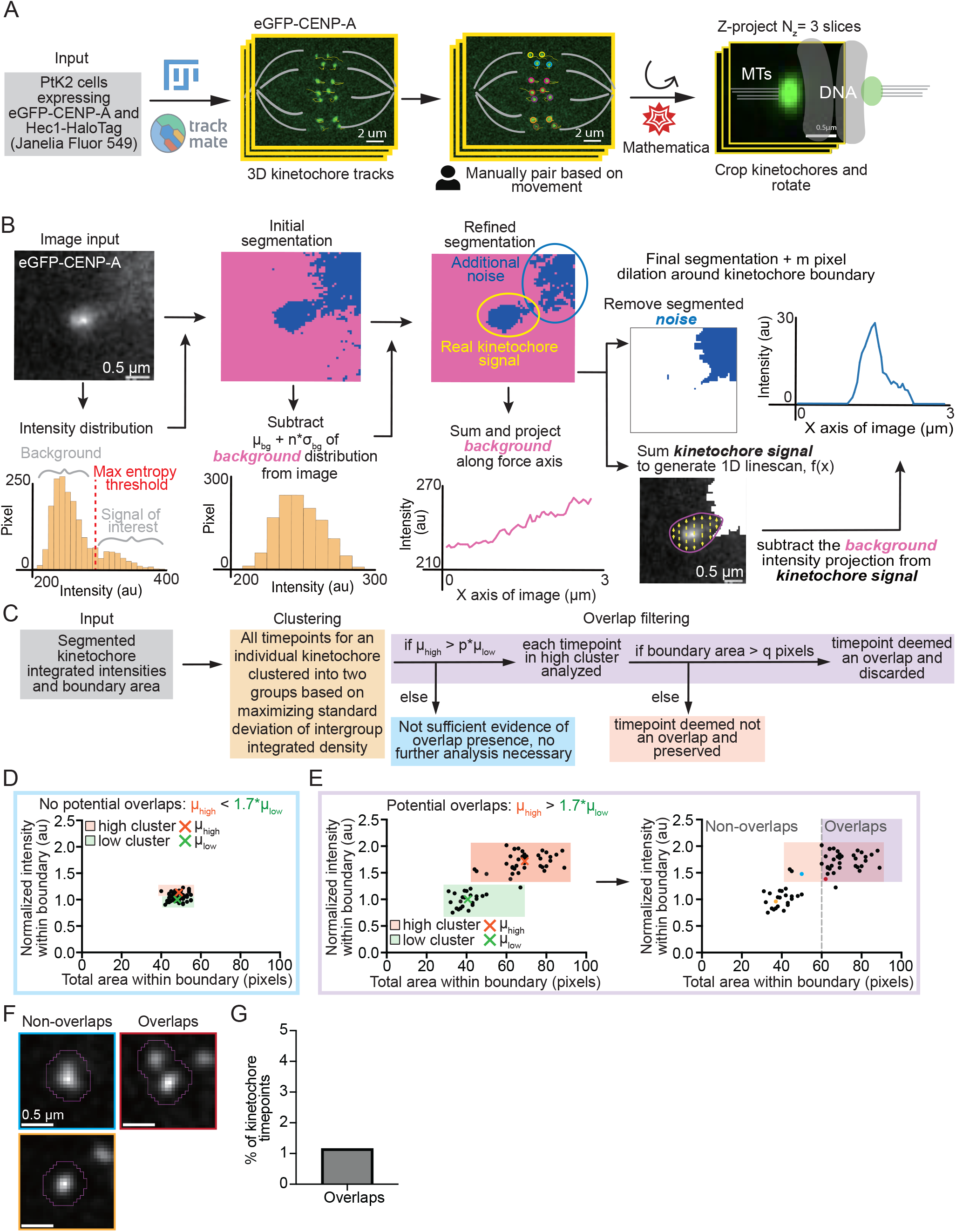
Image processing steps to extract kinetochore intensity signals for morphological analysis. (A) Schematic of workflow for kinetochore tracking (TrackMate, FiJi icons), sister pair assignment (manual, person icon), and force axis alignment (Mathematica icon, red star). Cropped kinetochore stacks are then rotated in Mathematica so that the sister kinetochore-kinetochore axis aligns with the image x-axis. A z-projection of the central z-stack slices (yellow) (here, N_z_ = 3) of each kinetochore is generated for downstream morphology analysis. Microtubules (grey lines) and chromosomes (grey) are drawn to orient the reader, together with the sister kinetochore (green oval). In this example eGFP-CENP-A images are used. (B) Schematic of the pre-processing workflow to segment kinetochores from the background. First, the raw kinetochore image is initially segmented using a maximum entropy threshold, dividing signal into background and kinetochore signal of interest. Segmentation is then refined using a background-based threshold, defined as the background mean µ_bg_ + n\**σ*_bg_. Background intensity is then projected into an intensity linescan, outside the segmented kinetochore region. Final segmentation then includes dilation around the kinetochore boundary by m pixels (here, m = 3). Additional noise is then removed for the final segmented kinetochore to be summed to generate a 1D intensity profile f(x), with the previously projected background subtracted from this 1D linescan. (C) Schematic of the workflow to identify kinetochore timepoints with “contaminating” signals from a neighbor kinetochore. The inputs are the segmented kinetochore intensities and its boundary area (grey box). Clustering is then performed to determine whether the individual kinetochore undergoes overlap filtering (yellow box). If the average intensity of the high cluster µ_high_ > p*µ_low_, where µ_low_ is the average intensity of the low cluster and p is a picked parameter, then overlap filtering is performed for each timepoint in the high cluster, and must have a boundary area > q pixels to be deemed an overlap and filtered (purple box). If µ_high_ < p*µ_low_, then kinetochore timepoints do not under overlap filtering (blue box). If boundary area < q pixels, then kinetochore timepoint is also not an overlap (red box). (D) Example plot of a kinetochore and their timepoints of the intensity within the kinetochore boundary and the area within the boundary, where timepoints are clustered into a high and low cluster but does not have potential overlaps (since µ_high_ < 1.7*µ_low_, p = 1.7 as in (Tran et al., 2025)). Each point is a timeframe of a single kinetochore. (E) Example plots of a kinetochore and their timepoints of the intensity within the kinetochore boundary and the area within the boundary, where timepoints are clustered into a high and low cluster (left plot) and is flagged for overlap filtering (right plot, since µ_high_ > 1.7*µ_low_). Kinetochore timepoints that are in the high cluster and have a boundary area > 60 pixels are classified as overlaps (right plot, shaded purple area). Each point is a timeframe of a single kinetochore. Red “x” indicates the µ_high_ while the green “x” indicates the µ_low_. Purple colored quadrant indicates the timepoints to be filtered, with a red colored dot indicating the example as in (F). Blue and yellow dot indicates non-overlapped timepoints as in (F). Dashed grey line indicates the total area threshold q. (F) Examples of eGFP-CENP-A kinetochore timeframes that are non-overlaps in the first column (blue and yellow boxes) and a flagged overlap in the second column (red box). (G) Plot of the percentage of kinetochore timeframes that are flagged and filtered out for potential kinetochore overlap from the control live imaged kinetochore dataset (1.2%; 76/6578 timepoints across m = 14 cells and n = 124 kinetochores) from our first application (Tran et al. 2025).

**Figure 3:**
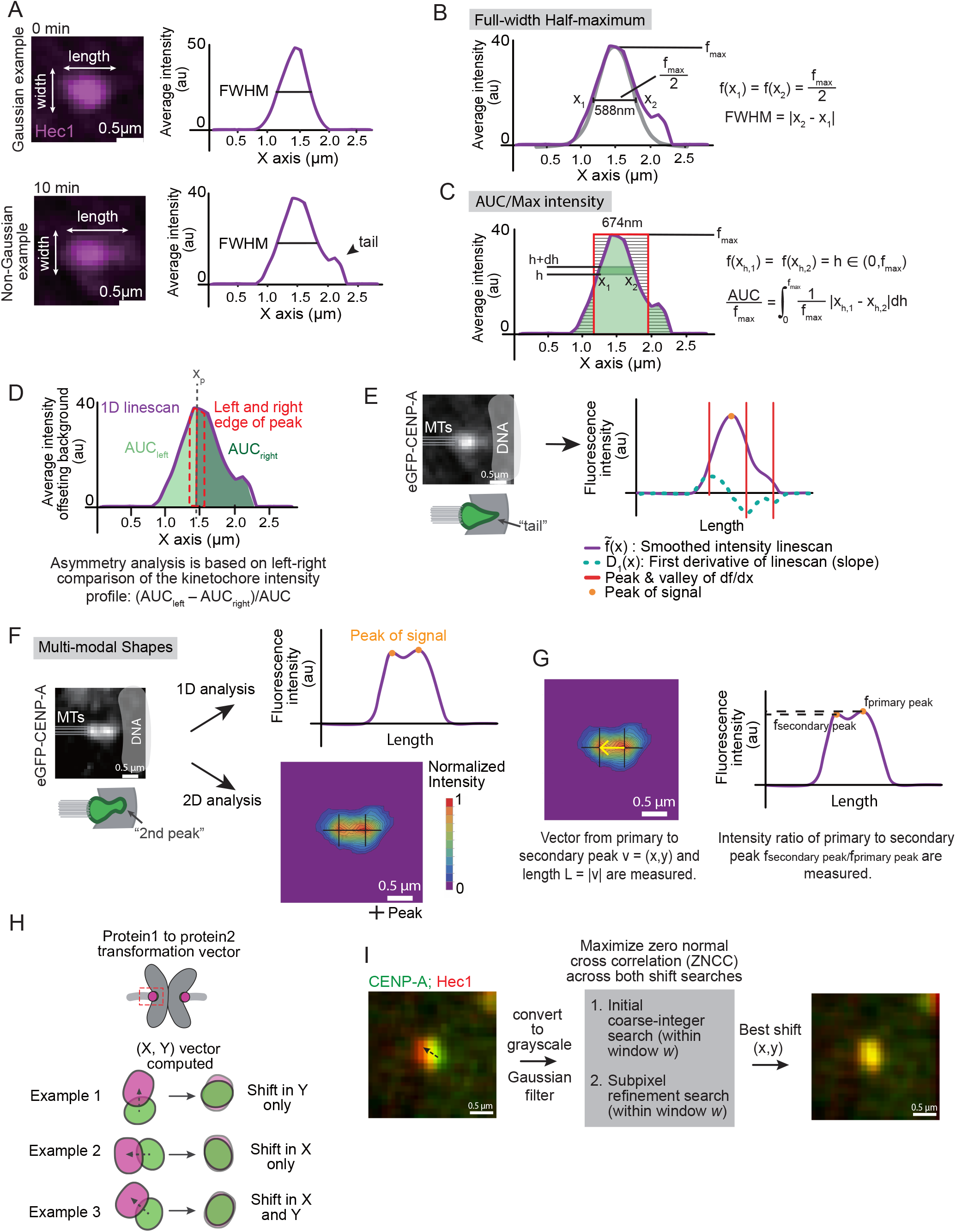
Developing metrics to define complex, non-Gaussian kinetochore morphologies. (A) Top image is a near Gaussian distributed example of a Hec1-HaloTag with JF549 dye distribution with corresponding 1D intensity linescan (purple) and the FWHM (black line) as kinetochore length metric. Bottom image is a non-Gaussian distributed example of the same kinetochore (10 min later) Hec1-HaloTag with JF549 dye distribution with corresponding 1D intensity linescan (purple) and the FWHM (black line) as kinetochore length metric. Black arrowhead indicates an intensity distribution change that occurs below the detection threshold of FWHM (black horizontal line). (B) FWHM length measurement of non-Gaussian kinetochore from (A) (bottom image). Purple line is the 1D intensity linescan and horizontal black line denotes the FWHM measurement (588 nm in this example). Equation for FWHM is shown on the right with f(x) the intensity at position x, and x_1_ and x_2_ the two positions where the half-max intensity is reached. (C) AUC/f_max_ length measurement of non-Gaussian kinetochore from (A) (bottom image). Purple line is the 1D intensity linescan, green shaded region indicates the area under the curve (AUC), f_max_ indicating the maximum intensity, black lines indicate the heights (h) of horizontal intensity slices to integrate, and the red box is equivalent to the AUC/f_max_ length metric (647 nm in this example). (D) Kinetochore intensity asymmetry analysis: Division of kinetochore intensity linescan (purple line) shown in (A) into a left and right side of the peak position x_p_. To score as asymmetric intensity, AUC calculated for the left (AUC_left_, light green area) and right (AUC_right_, dark green area) side must be consistent upon using the equation (AUC_left_ – AUC_right_)/AUC when the intensity linescan is divided into left and right sides at the peak, at the left edge, and at the right edge of the peak pixel (dotted red line area), to account for potential inaccuracy of peak position. (E) Kinetochore intensity tail analysis: Image of a kinetochore (eGFP-CENP-A) with a cartoon depiction of the “tail” formation. Corresponding smoothed 1D intensity linescan 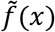 (purple), its first order derivative D_1_(x) (dotted blue), local extrema of D_1_(x) (red vertical lines), and primary peak of signal (yellow dot). (F) Kinetochore multimodality analysis: An image of a kinetochore (eGFP-CENP-A) is shown with a cartoon depiction of multimodality (multiple peaks, two in this example). A 1D and 2D peak detection analysis is used for each kinetochore timeframe. The 1D analysis is based on the same smoothed intensity projection linescan as in (E) (purple line) to detect peak formation along the X or Y axis, with identified peaks marked with yellow dots. The 2D analysis is based on intensity of signals across a 2D image above designated thresholds (bottom multicolored image), with identified peaks marked with “+”. (G) Schematic indicating additional peak measurements. Example kinetochore image (eGFP-CENP-A; left) showing distance between the two main peaks (denoted as |**v**(x,y)|) and indicated on image with black crosshairs with the vector shown (yellow arrow). Example smoothed 1D intensity distribution, 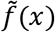, and the intensity ratio of the two peaks 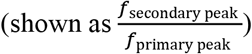. (H) Cartoon schematic of kinetochore protein1 (CENP-A, green) to protein2 (Hec1, magenta) with three examples of offset distributions where an (X,Y) transformation vector is computed to determine the distance with most overlap between distributions. Example 1 indicates a shift only in the Y axis, example 2 indicates a shift in only the X axis, and example 3 indicates a shift in both the X and Y axis. (I) Schematic of double channel analysis to define vector between kinetochore protein1 and protein2. A kinetochore image with double channels is shown (protein1 is eGFP-CENP-A in green, protein2 is Hec1-HaloTag with JF549 dye in red) that becomes converted to grayscale, smoothed with a Gaussian filter, undergoes an initial coarse-integer search method within a bounded window (denoted by *w*), then a second subpixel refinement search within bounded window *w*, with maximization of the zero normalized cross correlation (ZNCC) across both searches to obtain the vector (X,Y) (black dashed arrow, leftmost image) linking protein1 and protein2 that results in the highest overlap (overlapped CENP-A and Hec1, rightmost image).

**Figure 4:**
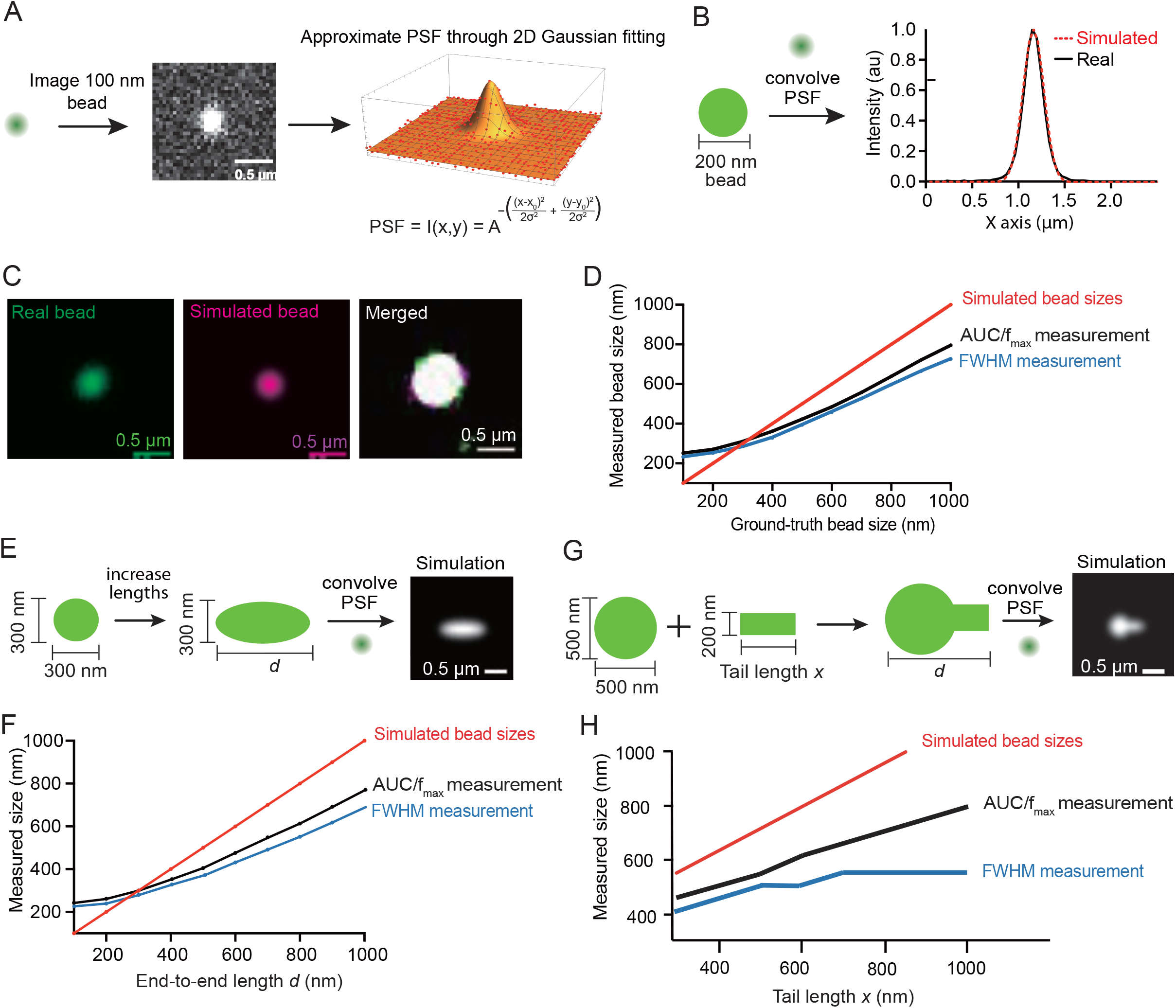
Validating the pipeline using simulations of ground-truth objects. (A) Schematic of the empirical measurement of the microscope point spread function (PSF). Fluorescent 100 nm beads were imaged and bead intensity was fit with a 2D Gaussian to generate an approximate PSF model. (B) Schematic of a ground-truth 200 nm bead convolved with the empirically measured PSF to generate a 1D intensity linescan for the simulated object (red dotted line) to be overlaid on a 1D intensity linescan of an imaged 200 nm bead (solid black line). (C) Representative images of iSIM-imaged 200 nm bead (green), a PSF-convolved simulated 200 nm bead (magenta), then a merged image (oversaturated to display the boundary). (D) Performance benchmark on simulated beads: measured bead size versus ground-truth bead size. AUC/f_max_ size estimate (black line), FWHM measurement (blue line), and the true simulated bead sizes are shown for a range of bead sizes. (E) Schematic of generating a library of uniformly stretched objects, increasing the end-to-end length *d*, and convolving with the empirically measured PSF to simulate an image. (F) Performance benchmark on uniformly stretched objects: ground-truth object lengths (red line), AUC/f_max_ size estimate (black line), and FWHM measurements (blue line) are shown. (G) Schematic of generating a library of “tailed” objects with an addition of a segmented rod to then extend by increasing the tail length *x* and convolving with the empirically measured PSF to simulate an image. (H) Performance benchmark on “tailed” objects with increasingly extended “tails”: ground-truth object lengths (red line), AUC/f_max_ size estimate (black line), and FWHM measurements (blue line) are shown.

We list in Table 1 all definitions of key variables that conceptually useful for developing the pipeline and important parameters in the code that can, and in some cases should, be user-adjusted according to the nature of the experimental data being analyzed. Table 1 includes variable names, definitions, default values from our first application dataset (Tran et al. 2025) and a brief description of what to consider when choosing values for each variable.

**Table 1:**
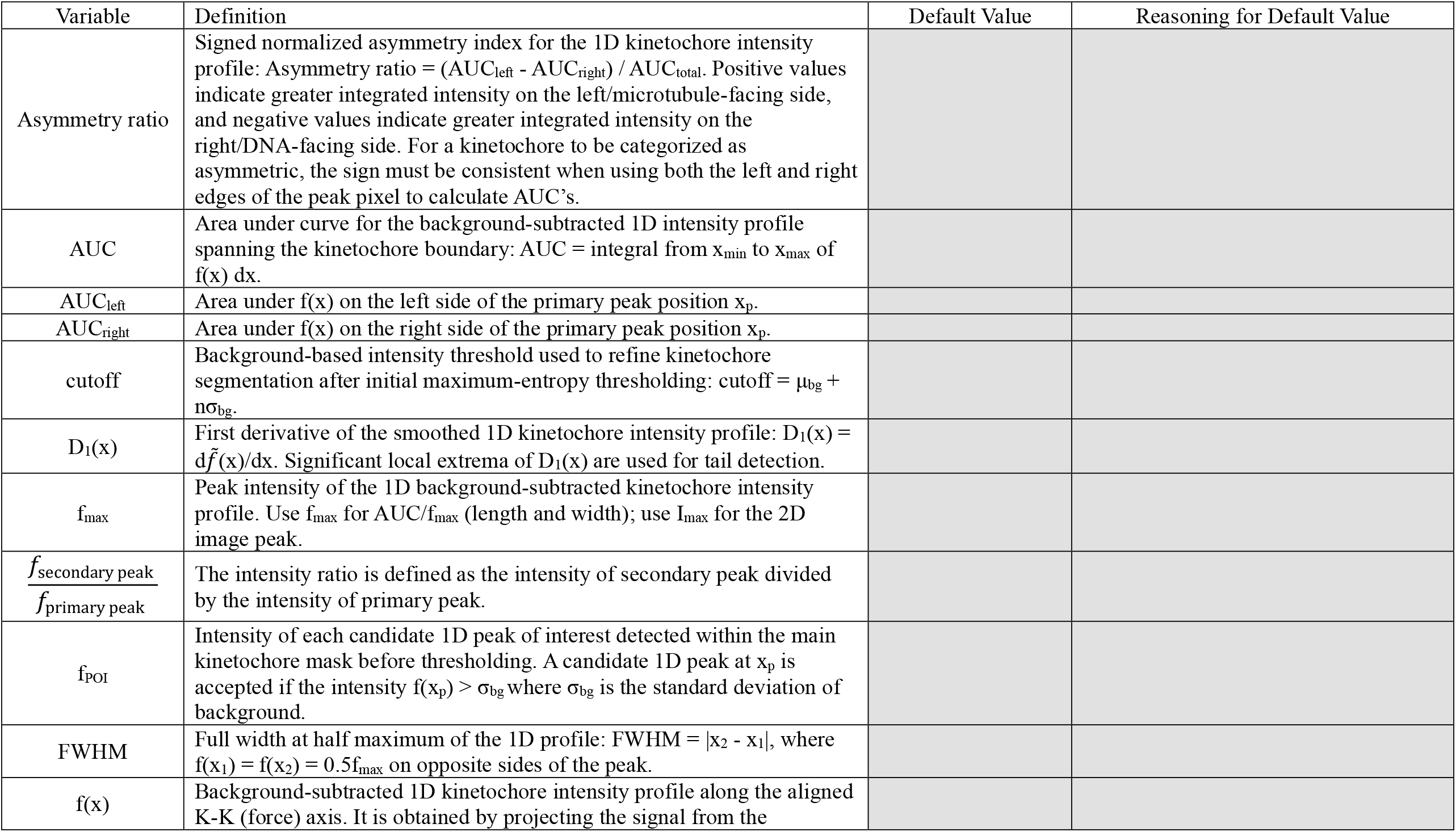
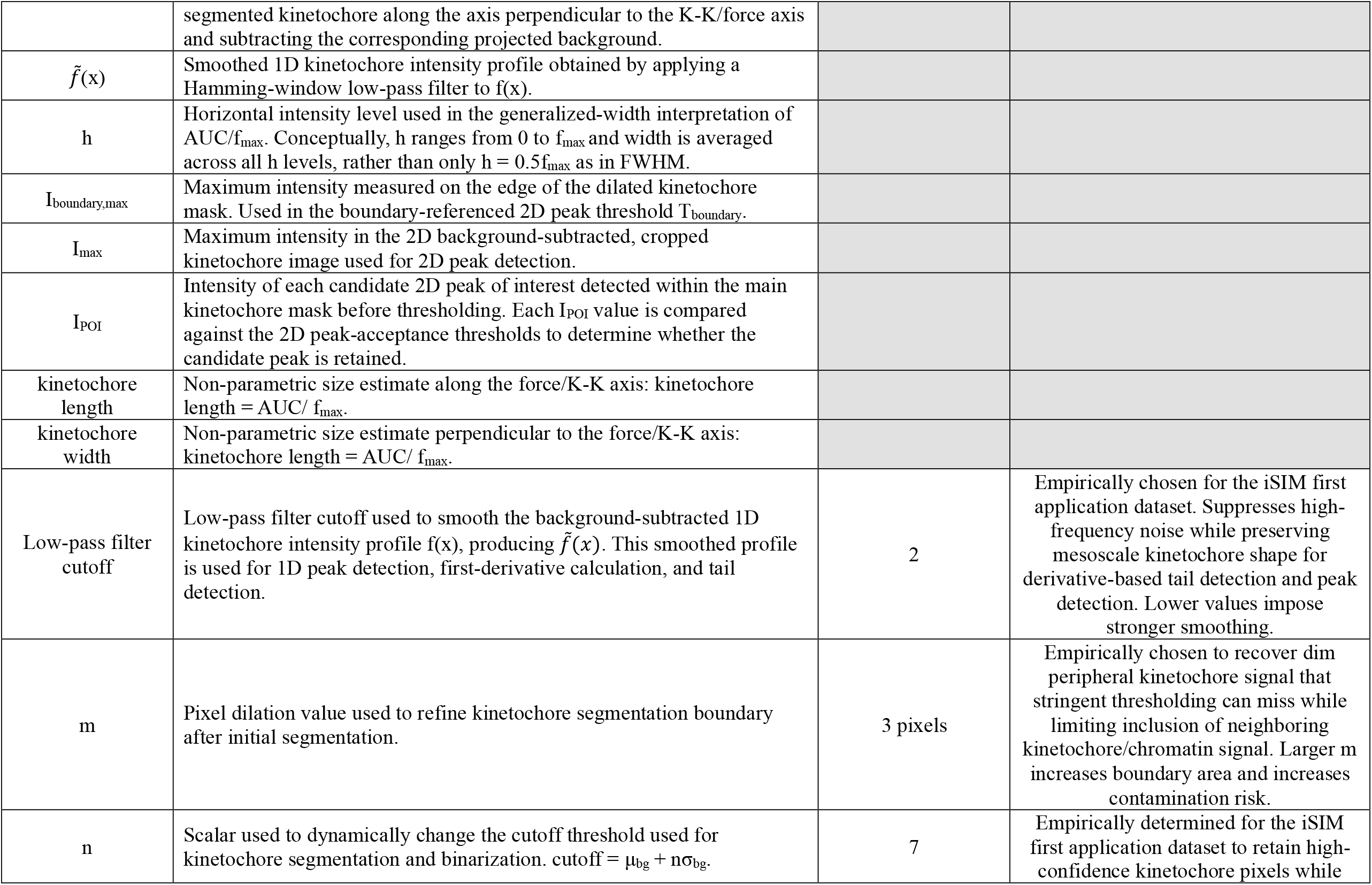
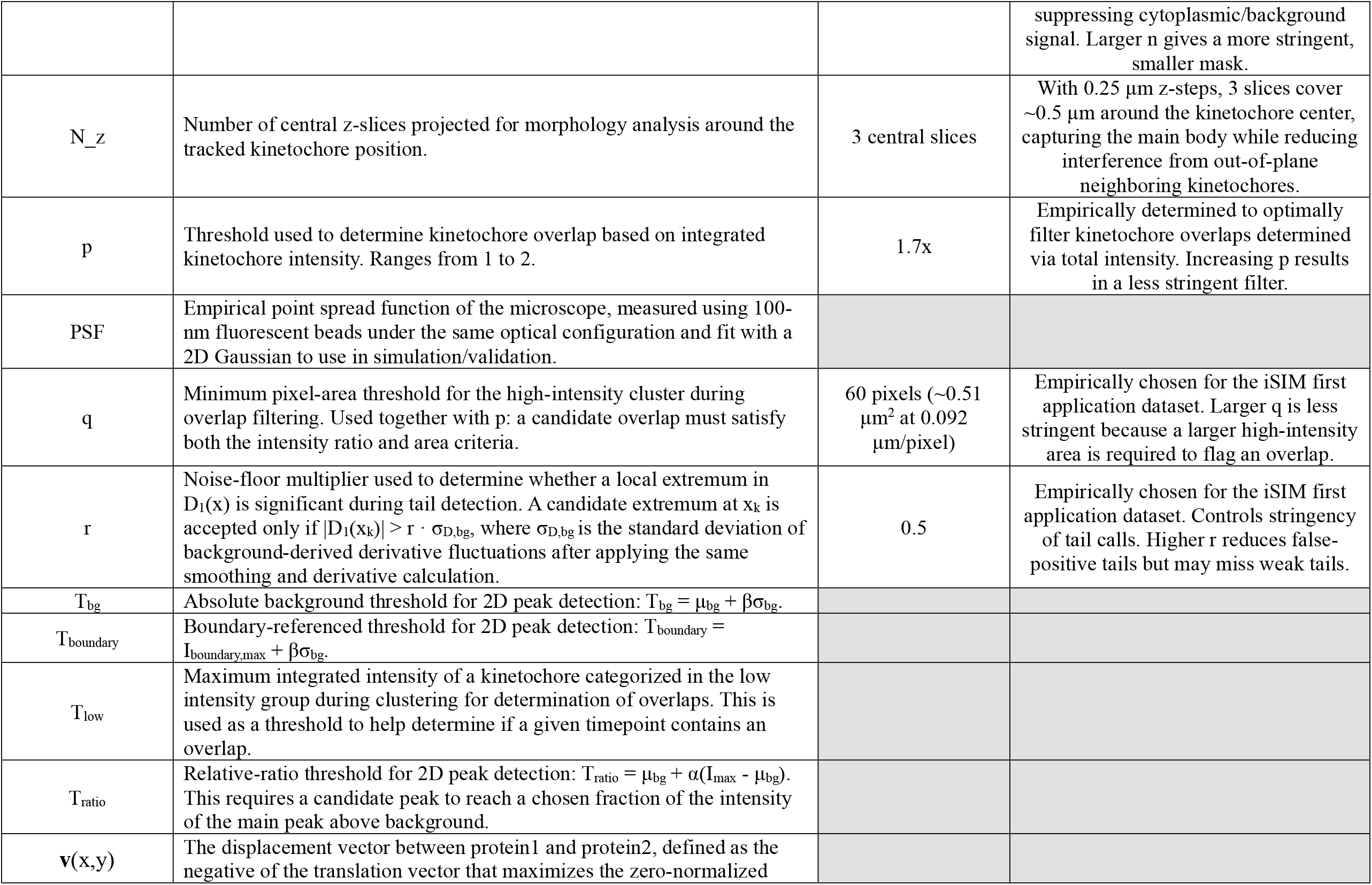
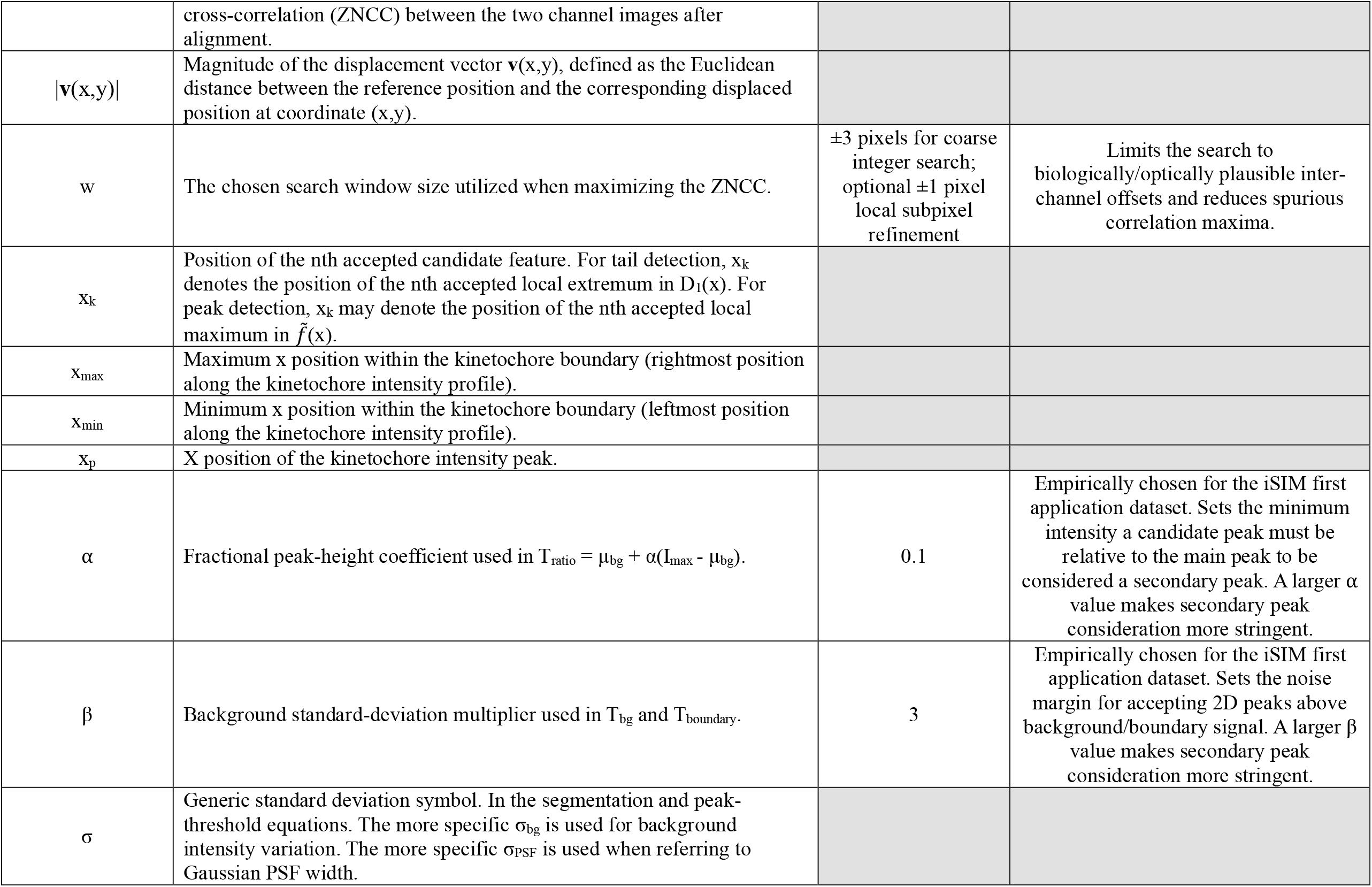
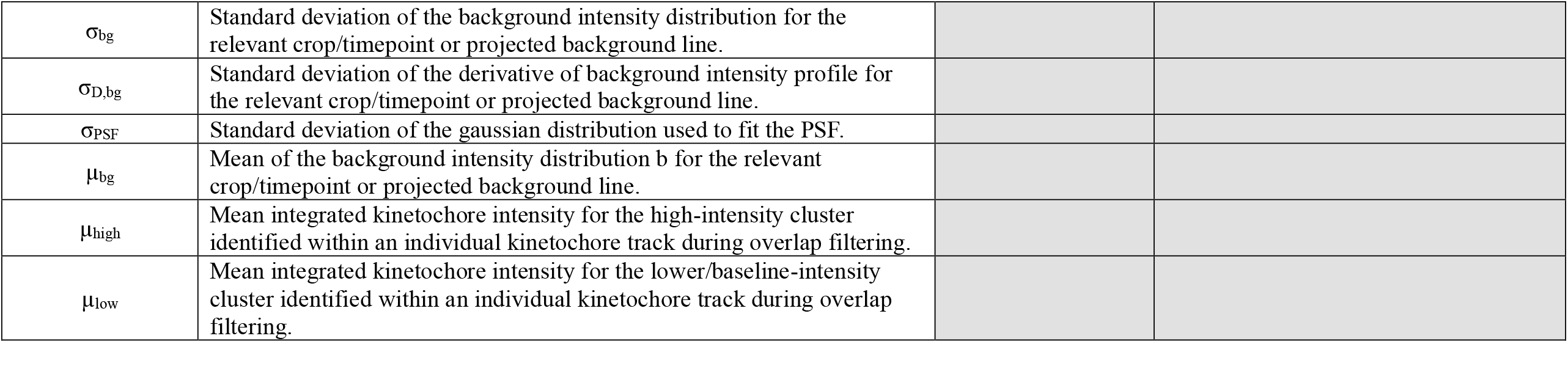
All variables, notations and measures including their definitions with default values, and reasoning for chosen default values (where applicable). Presented in alphabetical order.

### Overview of the computational pipeline for defining complex kinetochore morphologies from live cell movies

Figure 1 presents an overview of the computational pipeline we developed to quantify complex kinetochore morphologies, from input data to resulting output metrics. As input data (Figure 1, grey box), the pipeline accepts 4D movies of mitosis (timelapse movie with z-stacks) with a kinetochore protein fluorescently tagged. Ideal as input data to define complex kinetochore morphological changes over time are timelapse movies from microscopes offering high spatial resolution and limited photobleaching, such as from instant structured illumination microscopy (iSIM (York et al., 2013)). If the interest in defining kinetochore morphology is to learn about kinetochore structure, then kinetochore structural proteins should be chosen for fluorescent tagging. Input data can also include two kinetochore channels, namely two kinetochore proteins (protein1 and protein2) tagged in two different colors, which would provide information on the relative orientation of the intensity distributions of both kinetochore proteins.

The first step in the pipeline is to track kinetochores in 4D (3D space and time) during mitosis, which is done using TrackMate in FiJi (Figure 1, blue box) (Tinevez et al., 2017). These 3D tracks are then manually paired into sister kinetochore pairs (Figure 1, green box 1). The pipeline then relies on Mathematica (Wolfram Research, Inc., Version 14.3) for all further steps (Figure 1, red box) and outputs (Figure 1, purple boxes).

Image processing: To extract kinetochore intensity signals for morphological analysis (Figure 2), the pipeline rotates kinetochores to put them all along their force axis (kinetochore-kinetochore axis), crops kinetochores, projects 3D kinetochores to 2D then segments kinetochores while preserving as much of their low intensity shapes and generates the 1D projection which will be used for analysis(1^st^ purple box).

Morphological analysis: The pipeline then performs shape analysis, returning output metrics reflecting kinetochore dimensions, asymmetry, low intensity “tails”, number of intensity peaks (modality) and a 2D vector from kinetochore protein1 to protein2 (2^nd^ purple box). All along the way, the pipeline provides inspection points for visual quality control of segmentation and all shape analysis metrics (2^nd^ purple box): it displays the cropped kinetochore image, its segmentation, 1D intensity linescan, and intermediate analysis metric visuals.

Output compilation: In this final step, the pipeline outputs: (i) a master table with all cell IDs, kinetochore IDs (k_1_, k_2_, etc.), timepoints (t_1_, t_2_, etc.), and shape metrics; (ii) identification of which kinetochore timepoints are identified as “contaminated” with intensity from overlapping neighbor kinetochores, flagged for removal; (iii) movement analysis for each kinetochore and sister pair (velocity, standard deviation of position, kinetochore-kinetochore distance, etc.), in the event the user wants to correlate kinetochore morphological features with coincident movement features (3^rd^ purple box). Given the many existing computational pipelines for extracting kinetochore movement features, we do not focus on this aspect in the description of our pipeline here. While the pipeline we present here focuses on morphological analysis of a single kinetochore image, the pipeline is also able to extract how a single kinetochore’s morphology changes across time and of how morphological parameters are distributed across many kinetochores within a single cell or across different cells (Tran et al., 2025).

### Image processing steps to extract kinetochore intensity signals for morphological analysis

The first step is to convert raw, live-cell movies into clean, analyzable kinetochore signals. As a first application and demonstration (Tran et al., 2025), we imaged rat kangaroo (PtK2) cells expressing fluorescent markers (Hec1–Halotag with Janelia Fluor 549 dye and/or eGFP–CENP-A). In this case, cells were imaged in 4D using instant structured illumination microscopy (iSIM (York et al., 2013)) with 0.25 μm z-steps over 2.5 μm every 15 s. PtK2 cells remain relatively flat at mitosis which allows us to capture most kinetochores in limited z-stacks. Further, they have many fewer chromosomes (Lorenz & Ainsworth, 1972) than human cells which minimizes interference between neighbor kinetochore intensities. We obtained initial kinetochore trajectories with track stacks in Fiji using TrackMate (Fig. 1; Tracking). Although automated sister-pairing approaches are available (e.g., KiT (Armond et al., 2016; Harrison et al., 2022)), we performed manual sister-kinetochore pairing to ensure high-fidelity pairing, particularly under perturbative conditions where the metaphase plate and kinetochore oscillations are irregular (Tran et al., 2025); this step also serves as a verification that individual kinetochores are tracked correctly.

To analyze deformations in a force-relevant coordinate system, we cropped 3D kinetochore stacks to N_z_ slices and rotated them such that the kinetochore–kinetochore (K–K) axis aligns with the image X axis (with the Y axis being the metaphase plate axis and the Z axis perpendicular to the coverslip), using the K–K axis as a proxy for the dominant microtubule-generated force vector (Figure 1 and Figure 2A). Based on our first application (Tran et al., 2025) we chose a default N_z_ = 3 to capture the main kinetochore-of-interest’s signal while reducing out-of-plane overlapping kinetochores’ signal. From here on, we define kinetochore orientation such that pole-side microtubules are on the left (-X direction) and centromeric DNA is on the right (+X direction) (Figure 2A, rightmost image).

Defining the boundary of a diffraction-limited spot such as the kinetochore is inherently ambiguous. To define the boundaries of the kinetochore object, we used a two-step high-stringency segmentation strategy and then recovered the morphological boundary by dilation (Figure 2B). The premise is that a two-step segmentation strategy improves separation of the target kinetochore from nearby background signals while preserving more structural detail than a simple one-step thresholding approach. As a first step, we projected the central N_z_ Z-slices encompassing the detected kinetochore location. In our demonstrating examples, we use N_z_ = 3 (0.5µm). We selected this Z-range to capture the main body of the kinetochore while minimizing signal interference from other kinetochores. As a second step, we applied maximum entropy thresholding (Kapur et al., 1985), then elevating the binarization cutoff to a higher background criterion to aggressively suppress cytoplasmic background by following: With the mean of the initial segmentation background intensity defined as µ_bg_ and n multiplied by the standard deviation of the background (σ_bg_), we define the cutoff for refined segmentation as cutoff = µ_bg_ + n*σ_bg_. In our experimental and imaging demonstration conditions (Tran et al., 2025), we set the default n to 7 based on our empirical tests. When applying to the actual movies, this two-step segmentation method performed better than other established segmentation methods, such as testing Otsu’s method for thresholding (Figure S1A) as it improves the separation of nearby background kinetochore signals and retains only high-confidence signal pixels. If two or more disjointed components were detected, we selected the component that covers the center of the window or has the shortest distance to its center (Figure 2B; “refined segmentation”).

To recover the full morphological footprint—including dimmer peripheral signal that may be lost under stringent thresholding but is essential to shape definition—we dilated the resulting mask (e.g., by m pixels, but avoiding any other high intensity components) to capture the kinetochore “boundary,” and further restricted the analysis to this cropped kinetochore region to prevent “contamination” interference from nearby kinetochores or chromatin (e.g. in the case of a CENP-A label which can sometimes be present on the full chromosome). We empirically chose a default m = 3 pixels based on qualitative assessment when applied to our first application (Tran et al. 2025). Smaller dilations did not consistently capture the entire kinetochore signal while larger dilations increased the likelihood of capturing neighboring kinetochores’ signal or unnecessary background noise (Figure S1B). While the optimal number m of pixels dilation will depend on experiment-specific parameters (e.g. sample, imaging modality, camera), the principle remains the same. Finally, we projected the masked signal onto the K–K axis (that is oriented to be the X-axis) by averaging across the Y-axis and subtracting the corresponding projected background to obtain a one-dimensional, kinetochore-associated fluorescence profile suitable for quantitative analysis (Figure 2B, rightmost image).

To mitigate a major source of error, overlap between neighboring KTs, we implemented an overlap filter on each individual kinetochore timepoint (Figure 2C). By clustering images based on total intensity in the segmented area, we introduced an algorithm that flags kinetochores that show a significant change during the movie as determined by a combination of kinetochore intensity and measured pixel area. First, clustering of an individual kinetochore’s timepoints over its imaging lifetime into two groups is imposed, maximizing the standard deviation of the intergroup integrated density (Figure 2C, yellow box). Here, if the mean of the higher intensity group, µ_*high*_, is less than *p* * µ_*low*_, where *p* ∈ (1,2) is the preset threshold default as 1.7 as we experimentally found from testing on actual movies, then there is not sufficient evidence of overlap presence (Figure 2C, blue box; Figure 2D, left plot). If µ_*high*_ > *p* * µ_*low*_, then each timepoint within that kinetochore is further analyzed for overlap filtering and must meet two conditions as follows: (i) have a greater integrated intensity than T_low_, a threshold defined as the maximum intensity timepoint of the lower intensity and (ii) the boundary pixel area of the kinetochore at that timepoint is greater than a set pixel area, defined as *q* (default *q* = 60 pixels which equals ∼0.5um^2^ based on (Tran et al., 2025)) (Figure 2C, red box; Figure 2D, right plot). These thresholds (*p, q*) were validated empirically for our first application (Tran et al. 2025) dataset (Figure 2, D and E), and should be tuned to accurately capture and exclude overlapping kinetochores across other datasets where experimental parameters differ. Kinetochore timepoints that fulfill these conditions are then flagged as an overlap and discarded, and in our first application dataset (Tran et al., 2025) only 1.2% of timepoints are excluded from the analysis results (Figure 2F). This strategy is based on the observation that overlapping kinetochore signals produce substantially higher total intensity, while photobleaching in our iSIM movies is minimal. Because kinetochore brightness varies across cells and across individual kinetochores, clustering frames within each kinetochore track provides greater flexibility than applying a single absolute intensity threshold to all kinetochores. At the same time, requiring the high-intensity cluster to exceed a minimum segmented pixel area helps reduce false positives caused by transient intensity changes, such as kinetochores moving partially in and out of focus. Together, the intensity-based clustering and pixel-area requirements (AND gate) compensate for each other’s limitations and provide a more robust way to identify frames affected by neighboring kinetochore overlap than either of them could alone. In sum, the above image processing steps provide us with kinetochores that are aligned along the force axis, oriented with respect to the K-K axis, segmented away from background while retaining dimmer features, and largely reduce overlapping neighbor kinetochore signals.

### Developing metrics to define complex, non-Gaussian shape changes

#### (i)Non-parametric measure for kinetochore length and width: integrating the full intensity profile without a prior shape

Kinetochore fluorescence signals in many species, including most mammals, are near the diffraction limit of light. The simplest metric of kinetochore morphology is size. Here we define kinetochore length as size along the force (K-K) axis (X axis), and kinetochore width as size along the perpendicular axis in the XY plane (Y axis). The pipeline returns both length and width for each kinetochore from live-cell imaging, both determined as described here. The apparent “size” of a kinetochore is often summarized by the full width at half-maximum (FWHM) of a 1D intensity linescan profile (Magidson et al., 2015; Renda et al., 2020; Suzuki et al., 2014; Zhou et al., 2025). While FWHM is convenient and interpretable for approximately Gaussian, unimodal profiles (Figure 3A, top image and profile), it fails to fully capture complex, non-Gaussian kinetochore morphologies (Figure 3A, bottom image and profile). In particular, the FWHM is determined by a single intensity “slice” at 50% of the peak amplitude and therefore does not account for intensity changes below this threshold (Figure 3B). This creates a rigid, implicit assumption that the underlying kinetochore structure deforms by uniform broadening, making the FWHM effectively insensitive (“blind”) to biologically meaningful features that occur below the half-maximum threshold— such as low-intensity tails, multiple peaks, or asymmetries. We sought to overcome these shortfalls by developing a non-parametric measure of kinetochore length that integrates information across the full dynamic range of the signal and require minimal shape assumptions (Figure 3C). Because kinetochore assemblies are flexible protein complexes whose deformations need not conform to unimodal Gaussian shapes (Cimini et al., 2001; DeLuca et al., 2006; Magidson et al., 2015; Renda et al., 2020; Sacristan et al., 2024) we hypothesize that this non-parametric measure for morphological analysis will better capture “true” kinetochore dimensions and shapes, especially when their intensity profiles are non-Gaussian (Figure 3C). To capture a more accurate shape of non-uniform and non-Gaussian kinetochore intensity profiles, we define kinetochore length (and similarly for kinetochore width) as the total area under the curve (AUC) of the 1D kinetochore intensity profile divided by the peak intensity (*f*_*max*_) of that same profile 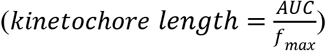. AUC was calculated as the integral of the smoothed, background-subtracted 1D kinetochore intensity linescan profile, 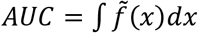, over the full x-axis projection range. The peak intensity, *f*_*max*_, was defined as the intensity of the primary peak in the same smoothed 1D profile. The presented relationship is equivalent to a generalized FWHM that utilizes the full width at all levels of the peak amplitude range from 0% to 100%, as opposed to only 50%, in an effort to more accurately capture kinetochore length while avoiding sampling bias that is inherent to the conventional FWHM method (Figure 3C). For Gaussian distributions, this relationship yields highly similar length measurements (within 7%) compared to the conventional FWHM method (Supplementary text). However, unlike the FWHM method, the integration of information across all intensity levels better captures biologically relevant non-uniform changes in intensity profiles found below the half-maximum threshold. In practice, we interpret this generalized formula for kinetochore size as a more adaptable descriptor that takes into account both simpler, Gaussian and more complex intensity profiles without requiring a Gaussian fit or intensity threshold.

#### (iii) Morphological analysis

##### (a)Asymmetry: directional deformation along the force-relevant axis

While kinetochore lengths and widths are important and useful descriptors of kinetochore morphology, they are not sufficient descriptors of kinetochore shape and shape changes. One parameter of interest is kinetochore symmetry or asymmetry, namely if the distribution of kinetochore intensity is symmetrical along the length and force (X) axis (and similarly for the width (Y) axis). Identifying and quantifying asymmetries in kinetochore intensity profiles can provide further insight into kinetochore molecular-scale attachment and structure and how these may change as a result of attachment force or geometry.

To quantify whether kinetochore intensity is symmetric between the microtubule-binding interface and the DNA-binding interface sides, we analyze the same background-subtracted 1D intensity profiles (Figure 2B, rightmost linescan) along the standardized K-K axis (microtubule side on the left, DNA side on the right). First, we find the x-position of the primary peak location (x_p_). Second, using this position we compute the AUC for the profiles left 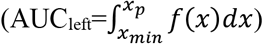 and right 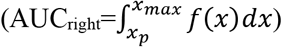 of the primary peak location (Figure 3D). Using these calculated areas, we define an asymmetry index as a signed normalized metric: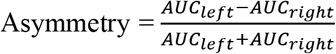. Because this approach relies on integrated area rather than a single threshold crossing, it remains informative for non-Gaussian profiles and supports direct comparison of deformation on the microtubule-facing side versus the DNA-facing side. For the qualitative “true/false” classification of asymmetry, we employ a flexible threshold by calculating the asymmetry twice: once using the left edge and again using the right edge of the x_p_ pixel (Figure 3D). For a kinetochore to be classified as asymmetric, it must demonstrate asymmetry when using both the left edge and the right edge of the x_p_ pixel and these asymmetries must possess the same sign, indicating a consistent asymmetry biased toward the DNA or microtubule side of the kinetochore. This left edge/right edge approach accounts for potential inaccuracies in identification of peak position. As a control, we calculated asymmetry for spherical beads with a diameter of 200 nm and found a 4.2% incidence of detected asymmetries, compared to a 16.5% incidence of detected asymmetries for our live-imaged first kinetochore application (Tran et al. 2025) dataset (Figure S2A). As a metric of the magnitude of asymmetry, we use (AUC_left_ – AUC_right_)/AUC, with 0 reflecting no asymmetry and 1 and -1 reflecting all intensity solely on the right or left side of the peak.

##### (b)Tail detection: automated classification of “tailed” versus “compact” profiles

Beyond continuous metrics of size and asymmetry, we implemented an automated classifier to detect kinetochore “tails” – extended, low-intensity appendages that are difficult to capture with FWHM-based measures but are qualitatively apparent. Automated detection of “tails” and the direction they extend (towards DNA or microtubules) can help quantitatively define orientation, frequency, and duration of kinetochore “tails”, all of which can in principle provide biologically useful insights on kinetochore structure and attachment status. While we focused tail analysis along the force and kinetochore length (X) axis, the same pipeline can be easily adapted to analyze tails along the kinetochore width (Y) axis.

To identify and classify “tail” structures, we analyzed the same background-subtracted 1D intensity linescan *f*(*x*) (Figure 3D, purple linescan). We used a Hamming-window low-pass filter to suppress high-frequency noise while preserving mesoscale shape, obtaining a smoothed kinetochore intensity profile 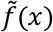 where we empirically determine how much to smooth the curve (Figure S2B, right column plots). Under-smoothing can result in detection of false “tails” (Figure S2B, top-middle plot) while over-smoothing can result in exclusion of true “tails” (Figure S2B, bottom-middle plot). Of note, this smoothing is dependent on the background noise levels that may differ based on the experimental context. We then computed the first derivative of 1D intensity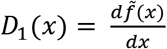 and identified significant local extrema (peaks/troughs) whose amplitudes exceed a preset noise floor *r* defined relative to the standard deviation of the background-derived derivative fluctuations, 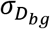 and 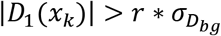. We define a “tail” as an extended, structured deviation from a single compact peak, where the first derivative displays at least three significant local extrema (Figure 3E). We use the first derivative of the 1D linescan because it provides a direct interpretation of extended slope structures andthis definition of the tail matches to our intuition a protrusion morphology.

The direction of the “tail” is determined by whether the first derivative has more than two local extrema and if more than one extrema is positive or negative. Specifically, a “tail” toward the left is specified by more than one positive extrema in the first derivative, while a “tail” extending toward the right is specified by more than one negative extrema. Here, we also quantified “tail” frequency of spherical beads as a control, and found 0.2% incidence compared to 5.7% in our live-imaged first application kinetochore dataset (Tran et al. 2025) (Figure S2C). This derivative-based method for “tail” detection can help distinguish between different deformation modes that kinetochores may exhibit, for example during merotelic attachments. This automated classification step provides an interpretable and objective morphology class for downstream analysis and hypothesis testing.

##### (c)Peak detection for multimodality

Next, we sought to capture and quantify kinetochore morphologies that deviate from a single compact peak during live-cell imaging, indicating the presence of a multimodal kinetochore structure which could have functional and biophysical consequences. To do this, we implemented peak-detection procedures to identify multimodal intensity distributions in both 1D and 2D (Figure 3F). We define multimodality as the presence of multiple local maxima, consistent with a multimodal signal from kinetochore substructures that cannot be well summarized by single-peak-width metrics. While we focused multimodal analysis along the force and kinetochore length (X) axis, the same pipeline can be easily adapted to analyze multimodality along the kinetochore width (Y) axis.

For 1D multimodality detection along the kinetochore length axis, we used the same smoothed 1D linescan intensity profile used for tail classification (Figure 3F, top). This allows for direct comparison between “tailed” and “multimodal” states and enables time-resolved tracking of identifying transitions between these two states. We identify a candidate peak-of-interest, *f*_POI_, as a local maxima that must exceed the standard deviation of the projected background intensity, σ_bg_, to be classified as a peak to avoid noise caused false positives.

While 1D projections provide a compact summary, they have the potential to merge distinct spatial features if peaks are separated off-axis or overlap in projection. To overcome this limitation, we also detect multimodality directly in the 2D image of the kinetochore to obtain higher spatial fidelity and sensitivity (Figure 3F, bottom). Starting from the background-subtracted, cropped kinetochore image, we detect local maxima in the 2D intensity field. Specifically, a 2D peak-of-interest (defined as *I*_*POI*_) was scored as a distinct peak if it met three criteria: (i) a relative ratio criterion requiring accepted peaks to be at least a fraction of the primary peak above background: *I*_*POI*_ > *T*_*ratio*_, where *T*_*ratio*_ = *α*(*I*_*max*_ − *μ*_*bg*_) + μ_*bg*_ in which α is the empirically chosen coefficient for the fraction of the main peak height (I_max_) and µ_bg_ is the mean background intensity; (ii) an absolute background criterion: *I*_*POI*_ > *T*_*bg*_, where T_bg_ = µ_bg_ +βσ_bg_ in which T_bg_ is the threshold over the background and β is empirically chosen multiplier for the background standard deviation to ensure accepted peaks are sufficiently brighter than background; (iii) a boundary-referenced criterion: *I*_*POI*_ > *T*_*boundary*_, where T_boundary_ = I_boundary,max_+ βσ_bg_ in which T_boundary_ is the kinetochore boundary-referenced threshold, I_boundary,max_ is the maximum intensity on the edge of the dilated kinetochore mask to ensure the peak-of-interest is beyond noise along the segmented boundary. The reason that there are more criterion for 2D than 1D is because they are more prone to the noise without projection. In our first application dataset (Tran et al., 2025), we set the default values of *α* = 0.1 and β = 3. If under-thresholded (Figure S2D, top row) or over-thresholded (Figure S2D, bottom row), false positives or false negatives may more likely to occur. If no peak satisfies these criteria, only the global maximum peak will be retained. A kinetochore is classified as 2D-multimodal when two or more significant peaks remain after filtering according to the above three criteria. Beyond detecting the number of kinetochore “peaks” and underlying substructures, relevant information about these peak intensities and distances between them can inform on how kinetochores deform and are organized. To this end, we quantify the distance between peaks in X and Y and also the intensity ratios that are background corrected of the second-to-first peak (Figure 3G). All together, the pipeline provides the number of kinetochore intensity peaks, and if more than one peak, the relative brightness of peaks and the distance between peaks.

##### (d)Double channel analysis to define vector between kinetochore protein1 and protein2

Lastly, we sought to determine the relative position of two different populations of live-imaged fluorescently-tagged kinetochore proteins within the same kinetochore, where protein1 and protein2 are labeled in different colors. Knowing this distance and direction provides information regarding the nanoscale architecture and organization of proteins within the kinetochore and how they may shift over time and in different attachment scenarios. To do this, we used movies containing both inner (protein1) and outer (protein2) kinetochore markers and quantified the vector between protein1-protein2 in various orientations using a 2D image registration approach (Figure 3H). After cropping individual kinetochores and rotating them so that the sister-kinetochore axis is aligned with the x-axis, matched image regions from the two channels were extracted. Each image was adjusted in intensity, converted to grayscale, and smoothed with a Gaussian filter. We first performed a coarse integer-pixel search within a bounded search window, w, to compute the initial transformation while constraining the search area (default value w = 3 pixels). When this integer is found, we then perform a second, subpixel refinement search using Nelder-Mead optimization to interpolate within 1 pixel to further compute the transformation vector. During both the coarse integer and subpixel refinement search, we identified the (X,Y) translation that maximized the zero-normalized cross-correlation (ZNCC) between the two channel images (Figure 3I). Finally, the displacement vector between the two channels was defined as the negative of the shift required to align protein1 to protein2 and was converted from pixels to nanometers using the known pixel size. Because the images were pre-aligned to the load-bearing axis, the X component of the displacement reflects relative separation along the force-bearing direction, whereas the Y component reflects lateral offset perpendicular to that axis. In our first application dataset (Tran et al., 2025), the relative distance between the Hec1 and CENP-A populations was 56.5 nm in X and 0.7 nm in Y (Tran et al., 2025). This is in line with the expected distance between the two protein populations (Roscioli et al., 2020; Wan et al., 2009), helping to validate image processing steps in our pipeline. Notably, the registration-based approach we present here does not require explicit Gaussian fitting of either signal and is therefore well suited to irregular or non-Gaussian kinetochore shapes.

### Compiling data

Figure 2 presented the pipeline’s image processing steps to segment, extract and prepare kinetochore intensity signals prior to morphological analysis. Figure 3 presented the pipeline’s methods and metrics for morphological analysis, focusing on kinetochore size (length, width), asymmetry, low intensity “tails” and multimodality, i.e. peak analysis. The next step in the pipeline is compilation of all above metrics for each kinetochore (k_1_, k_2_, etc.) at each timepoint (t_1_, t_2_, etc.), which is returned to the user in the form of an indexed master data table. The user can then use this table to ask and answer biological questions such as: how the size and shape of a kinetochore evolves during its lifetime at mitosis; how kinetochore size and shape metrics compare between kinetochores of the same cell or of different cells; how kinetochore size and shape correlate with sister kinetochore size and shape or with kinetochore movement.

### Validating the pipeline using simulations of ground-truth objects

In an effort to validate the above computational pipeline development in an imaging regime relevant to live kinetochores, we performed simulation-based validation of kinetochore size measurements that incorporates the empirically parameterized Gaussian point-spread function (PSF) of our microscope (in our first application, this was an iSIM system (Tran et al. 2025)). We designed a set of biologically possible known kinetochore morphologies – ranging from uniform stretching to directional extensions (“tails”) – and generated synthetic images by convolving these ground-truth shapes with the measured PSF. This approach allows direct testing of whether our pipeline can reliably capture continuous shape changes under diffraction-limited blurring and can be benchmarked against the ground truth data of defined geometries. Below we present our validation efforts, and we include the code for users to replicate this effort for their microscope and different object shapes as a separate module of this computational pipeline.

#### (i) Empirical PSF characterization of the iSIM system

We first measured the system PSF experimentally using 100 nm fluorescent beads imaged under the same optical configuration used for live cell imaging in our first application dataset (Tran et al. 2025). We then fit the bead intensity distribution to a 2D Gaussian model to obtain PSF parameters (σ_PSF_) (Cole et al., 2011; Zhang et al., 2007), which served as the empirically parameterized Gaussian PSF for subsequent simulations (Figure 4A). This empirical calibration ensures that the simulated image formed matches the microscope’s effective performance.

#### (ii) Convolution-based forward simulations of deformed kinetochores

Using the empirically parameterized Gaussian PSF, we generated synthetic images of kinetochore-like objects via forward image formation, comparing both the intensity distribution and images of an imaged 200 nm bead and a 200 nm simulated bead (Figure 4, B and C). Specifically, we created a library of ground-truth shapes representing simplified but biologically relevant morphologies: (i) symmetric “beads” with a range of known radii to model uniform kinetochore size changes (Figure 4D); (ii) ellipsoids/ovals to model anisotropic size changes (Figure 4, E and F); and (iii) asymmetric extensions appended to an otherwise compact core to model “tailed” structures (Figure 4, G and H). Simulated images were then passed through the same preprocessing and metric-extraction pipeline used for live movies (background subtraction, 1D projection, and computation of FWHM, AUC/f_max_, asymmetry, and “tails”), enabling direct evaluation of metric behavior under defined ground-truth morphologies.

#### (iii) Performance evaluation: linearity and sensitivity

We assessed the linearity and sensitivity of our non-parametric kinetochore length measurement equation using PSF-convolved simulated objects with known ground-truth geometries. We first simulated circular bead-like objects of increasing size to test performance on compact, near-Gaussian signals (Figure 4D). For these objects, measured kinetochore length scaled approximately linearly with ground-truth size and behaved similarly to FWHM, indicating that it preserves the expected size-tracking behavior for simple Gaussian-like objects (Figure 4D). We then tested uniformly elongated, oval objects to determine whether the non-parametric derived kinetochore length remained accurate for compact but non-circular shapes (Figure 4E). Similar to the bead simulations, measured kinetochore length and FWHM showed broadly comparable trends for these oval objects, suggesting that both metrics can report uniform elongation when the signal remains in one peak and is symmetric after PSF convolution (Figure 4F). The key difference emerged in the tail simulation, where a low-intensity extension was appended to a circular object (Figure 4G). In this case, the measured kinetochore length using the non-parametric method remained linearly responsive to increasing tail length because it integrates signal across the full intensity profile. In contrast, FWHM showed limited sensitivity when the added extension remained below the half-maximum threshold (Figure 4H). As expected, both measured size metrics (AUC/f_max_ and FWHM) plateau as object size decreases due to the diffraction limit (Figure 4D, F, H). Together, these simulations demonstrate that our pipeline (i) behaves as expected on Gaussian-like objects, (ii) preserves linear tracking of size changes, and (iii)substantially improves sensitivity to biologically relevant, low-intensity asymmetries that conventional FWHM-based quantifications are not designed to capture.

## Discussion

In this work, we present a computational pipeline to quantify kinetochore morphological changes in live cells. While much computational work has been done over the last two decades to define the relative nm-scale positions of layers of kinetochore protein species within the kinetochore (Wan et al., 2009), including in live cells (Dumont et al., 2012), much less work has been done to define the shapes of each of these layers. Challenges include the small, diffraction-limited size of kinetochores, kinetochore shapes often being complex, irregular and fast-changing, the limited signal-to-noise ratio of kinetochores in live images due to photobleaching and kinetochores being deep inside cells, and kinetochores moving rapidly at mitosis. Most work describing the shape of kinetochore layer intensity distributions relied on the FWHM metric to describe how wide the distribution of proteins was (Magidson et al., 2015; Renda et al., 2020; Suzuki et al., 2014; Zhou et al., 2025). Yet, we have long known that kinetochore layers can take on shapes that are far from Gaussian or other common descriptors (Cimini et al., 2001; DeLuca et al., 2006; Magidson et al., 2015; Sacristan et al., 2024). The pipeline we present fills this gap.

The pipeline we present integrates kinetochore tracking with sister pairing, rotation into a force-relevant coordinate system, segmentation, automated exclusion of kinetochores whose intensities overlap with neighbor intensities, and 1D intensity projection (Figures 1-2). The pipeline moves beyond Gaussian assumptions to quantify irregular shapes using a non-parametric size descriptors (AUC/f_Max_) of length and width that integrates the full intensity profile rather than relying on a single 50% isointensity slice as the FWHM metric does. We present classifiers that report on morphology – directional asymmetry, low-intensity “tails,” and multimodal peak structure – capturing morphological patterns that are not commonly reported by conventional quantitative metrics for kinetochore shape (Figure 3). For two-color movies, we incorporate a registration-based fitting approach to quantify the relative displacement between protein1 and protein2 layer signals, providing a vector-based readout of intra-kinetochore architecture that captures both separation along the force-bearing axis and lateral offset between layers (Figure 3, H and I). Finally, we establish that our size metric readouts reflect underlying structural changes by using simulations that convolve ground-truth objects with an empirically measured PSF (Figure 4). These validations demonstrate that AUC/f_max_ closely scales with real size for near-Gaussian objects while remaining sensitive to low intensity “tails” and complex shapes where FWHM fails.

One limitation of our pipeline stems from the complexity of potential kinetochore shape changes and the inherent ambiguity between signal and noise when dealing with small objects. Since measured kinetochore lengths and widths depend on the pattern and signal intensity, low intensity signal can lead to potential underestimation of the actual object size. However, we find that the AUC/f_max_ remains robust to detecting relative changes in object size (Figure 4F). This allows us to infer what shape changes occur under various contexts. Finally, we note that the shape classifiers we present in this work (asymmetry, tails, and multimodal intensity) will benefit from being further refined, for example by developing a machine learning-based classifier for kinetochore shapes.

In summary, our computational pipeline presents a flexible and robust kinetochore size measurement method and brings forward new metrics for defining complex kinetochores morphologies. This approach improves on the limitations of traditional FWHM measurements and provides a more comprehensive description of kinetochore structure in live cells. The first test case of our pipeline was to mammalian metaphase kinetochores (Tran et al., 2025) and its application underscores its biological relevance to understanding the impact of attachment geometry, mechanical force, and molecular composition on complex features of kinetochore structure. Looking forward, we believe that this pipeline will be a useful tool to ask a broad range of questions ranging from kinetochore material properties and their molecular basis to live dynamics of attachment formation and error correction to the coordination of individual kinetochore-microtubule plus-end dynamics at a single kinetochore. More broadly, we believe that the pipeline we present may be useful for defining the live structural dynamics of other small subcellular structures that take on complex, irregular shapes.

## Acknowledgement

We thank Samuel J. Lord, Tanner Fadero, Nico Stuurman, Nigel J Burroughs, Wallace Marshall, and the Dumont lab for helpful discussions. This work was supported by NIH R35GM136420 (S.D.) and NSF Center for Cellular Construction DBI-1548297 (S.D.), NIH T32GM007810 (V.M.T), NIH T32GM139786 (V.M.T.), NIH T32EB009383 (V.M.T.), NIH F31GM156031 (V.M.T.), and the NSF Graduate Research Fellowship Program (C.J.R.).

## Author contribution

Conceptualization (J.T, V.M.T., S.D.); data curation (J.T., V.M.T); formal analysis (J.T.); funding acquisition (V.M.T, C.J.R., S.D.); investigation (J.T., V.M.T.); methodology (J.T., V.M.T., C.J.R., S.D); project administration (S.D.); resources (J.T., V.M.T); software (J.T.); supervision (V.M.T., S.D.); validation (J.T., V.M.T.); visualization (J.T., V.M.T., C.J.R., S.D.); writing – original draft (J.T.); writing – review and editing (J.T., V.M.T., C.J.R., S.D.).

## Supplemental Figure Legends

**Supplemental Figure 1:**
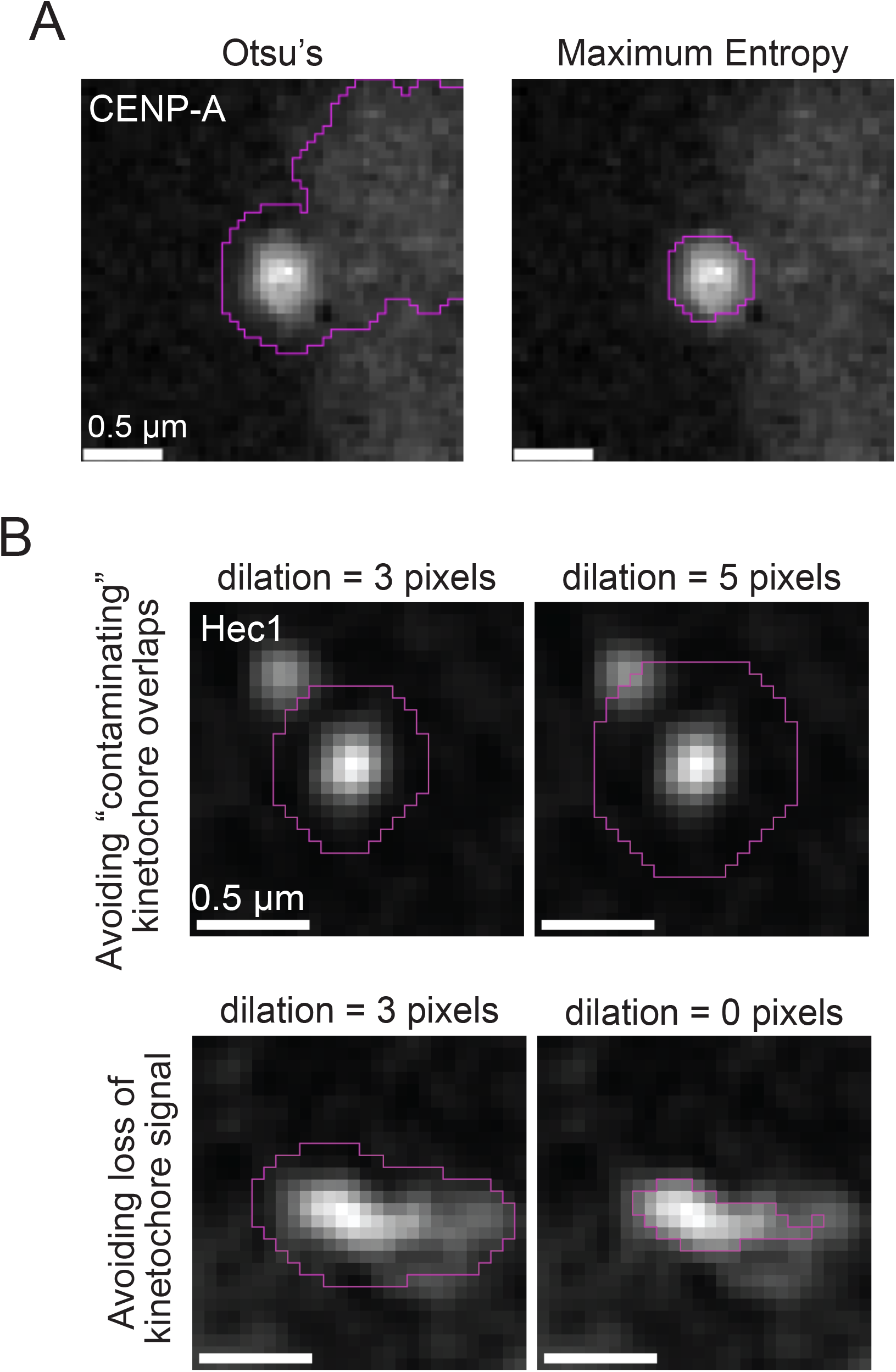
Validation of kinetochore image processing segmentation methods. (A) Example kinetochore images (eGFP-CENP-A) showing the segmented kinetochore signal including chromosome arm noise from Otsu’s thresholding (left) and the segmented kinetochore signal from the chosen maximum entropy thresholding (right). (B) Example kinetochore images (Hec1-HaloTag JF549 dye) showing comparison of increasing m pixel dilation that includes kinetochore noise (top row) and comparison of decreasing m pixel dilation that excludes kinetochore signal (bottom row).

**Supplemental Figure 2:**
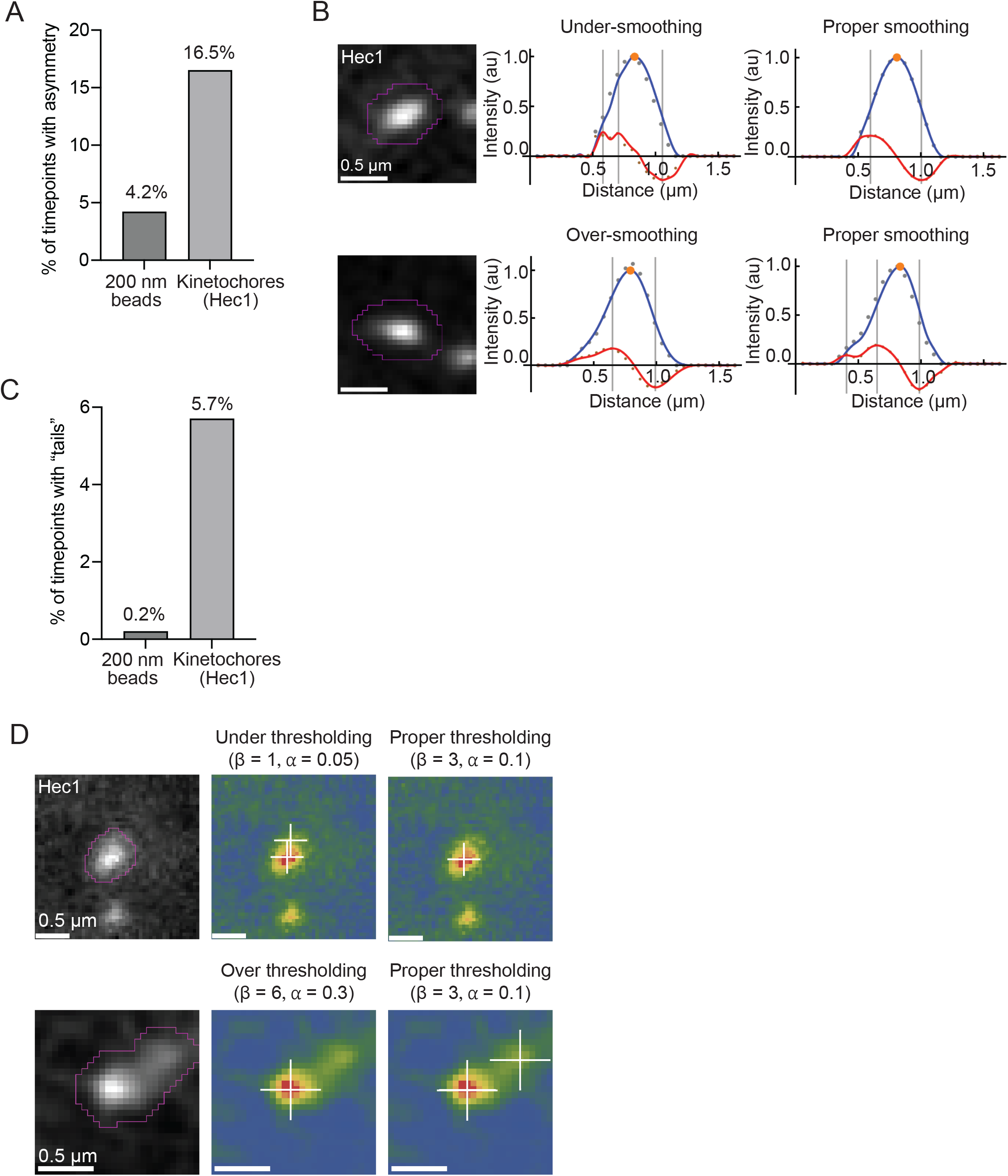
Validation of morphological analysis. (A) Plot of the percentage of timepoints with intensity asymmetry detected for imaged 200 nm beads (4.2%; m = 22 beads, n = 504 timepoints) and for a live-imaged Hec1-HaloTag with JF549 dye kinetochore dataset (16.5%; m = 124 kinetochores, n = 6578 timepoints) imaged in the same wavelength. (B) Example kinetochore images (Hec1-HaloTag with JF549 dye) showing comparisons between smoothing parameters. Top row shows a kinetochore image with corresponding 1D intensity linescans (blue lines), the main peak (yellow dots), and the first derivative curve of the linescans (red lines) when under-smoothed where low-pass filter cutoff = 3 (middle column) and properly smoothed where low-pass filter cutoff = 2 (right column). Bottom row shows a kinetochore image with corresponding 1D intensity linescans (blue lines), the main peak (yellow dots), and the first derivative curve of the linescans (red lines) when over-smoothed where low-pass filter cutoff = 1 (middle column) and properly smoothed where low-pass filter cutoff = 2 (right column). (C) Plot of the percentage of timepoints with “tails” detected for imaged 200 nm beads (0.2%; 22 beads, 504 timepoints) and for a live-imaged Hec1-HaloTag with JF549 dye kinetochore dataset (5.7%; 124 kinetochores, 6578 timepoints) imaged in the same wavelength. (D) Example kinetochore images (Hec1-HaloTag with JF549 dye) showing comparisons between 2D peak thresholding parameters. Top row shows a segmented kinetochore (left), when it is under-thresholded with β = 1, α = 0.05 (middle), and with proper thresholding with β = 3, α = 0.1 (right). Bottom row shows a segmented kinetochore (left), when it is over-thresholded with β = 6, α = 0.3 (middle), and with proper thresholding with β = 3, α = 0.1 (right). White crosshairs indicate the detected peaks.

## Materials and Methods

The computational pipeline we present can be used with different experimental contexts, including different cell lines, tagged kinetochore proteins, microscopes and image acquisition parameters. Below we describe the experimental work behind our first application dataset (Tran et al. 2025) that led to the example images and data provided herein (Figures 2 and 3 and Supplemental Figures 1 and 2).

### Mammalian cells

PtK2 cells stably expressing eGFP-CENP-A were transiently expressing Hec1-HaloTag with lentiviral infection as in Tran et al., 2025. All PtK2 cells were cultured at 37 °C and 5% CO_2_ in MEM (Invitrogen) supplemented with 5% sodium pyruvate (Invitrogen), 5% non-essential amino acids (Invitrogen), 5% penicillin/streptomycin (Invitrogen), and 10% qualified and heat-inactivated fetal bovine serum (Gibco). Cells were plated in 35 mm glass-bottom dishes (poly-D-lysine coated; MatTek Corporation) for live imaging experiments.

### Beads

Microspheric 100 and 200 nm TetraSpeck beads (Invitrogen) were pipetted (10-15 µL) into live-cell imaging dishes as plated above. Beads were left with cells in imaging dishes for at least 15 minutes to allow for settling onto glass bottom before imaging.

### Microscopy

Imaging was done using an inverted Nikon Ti-E equipped with a VT-iSIM super resolution module, 100-200 mW laser lines (405, 488, 561, and 642 nm), 100x 1.45 NA PlanApo oil objective (Nikon), Cairn Optospin emission filter wheel (ET 450/50m, ET 525/50m, ET 595/50m, ET 655lp, and ZET 405/488/651/640 m; Chroma Technology Corp.), and a Hamamatsu Quest interline-CCD camera. Cells were imaged with 11-13 z-planes 0.25 µm apart in 488 and 561 every 15 s, and a single central z-plane in 640 captured every 5 timepoints. After VT-iSIM imaging, images were deconvolved using Microvolution software (Microvolution). Images were collected at bin = 2 (92 nm/pixel) on Micro-Manager (2.0.0). Cells were imaged in a humidified stage-top incubation chamber (Okolab) at 30 °C with 5% CO_2_. For experiments imaging 100 nm beads for point spread function generation (described below), beads were imaged with 13 z-planes 0.25 µm apart in 488 and 561, adjusting laser power as needed to obtain similar signal-to-noise ratios for each channel imaged. For timelapse imaging of 200 nm beads to test morphological analysis, the same imaging parameters as used for live cells were also used for beads; beads were only imaged for 6 minutes.

### Point spread function (PSF) generation and use

For each bead, a two-dimensional Gaussian function

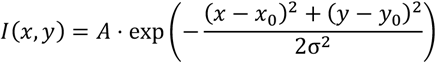

was fitted to the bead’s intensity profile in the central z-plane using custom routines in Mathematica. Here, A is the peak amplitude, (*x*_0_, *y*_0_) denotes the bead center, and σ represents the lateral standard deviation of the system’s PSF. The fitted σ was used to construct a representative PSF kernel. This kernel was then convolved with idealized shapes (e.g., beads, stretched ellipses, tails) to simulate diffraction-limited images for validation of length quantification.

### Computational pipeline

The Mathematica code behind the complete computational pipeline, including for validation and simulation, can be found here: https://github.com/Jinghui-Tao/kinetochore-morphology-pipeline.

## Supplementary Text

### Relating the AUC/*f*max measurement to the full-width-at-half-maximum measurement

We present a mathematical deduction supporting our application of a non-parametric measurement method for the morphological analysis of diffraction-limited, mammalian kinetochores during live-cell imaging. The goal of this section is to clarify why the AUC/*f*max measurement can be used as a generalized, non-parametric measure of kinetochore width and compare this measure to the conventional full-width-at-half-maximum measurement (FWHM). Specifically, we show that AUC/*f*max represents the average width of the normalized intensity profile across all intensity thresholds. This makes it sensitive to the full shape of the profile, including asymmetric tails and multi-peaked distributions, while still reducing to a fixed proportional relationship with the conventional FWHM measurement for an ideal Gaussian profile.

We begin with a non-negative one-dimensional kinetochore intensity profile.

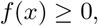

with peak intensity

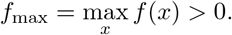

We first normalize the intensity profile against the peak intensity

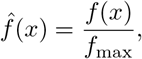

so that the intensity profile is bounded between 0 and 1

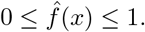

Since AUC is defined as the area under the 1D kinetochore intensity profile, AUC/*f*max measurement is then

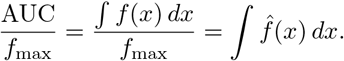

For each threshold h is defined as a given horizontal intensity level

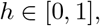

define the superlevel set

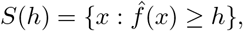

and let its one-dimensional measure, or length, be

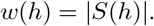

Here, *w*(*h*) is the width of the normalized profile at fractional intensity threshold *h*. In particular, the conventional full-width-at-half-maximum is the width at *h* = 0.5:

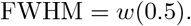

For any value 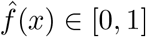, to be able to switch the integration to along h rather than x, we can write

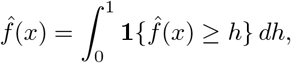

where **1**{·}is the indicator function. Integrating both sides over *x* and exchanging the order of integration gives

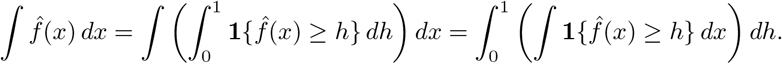

Because

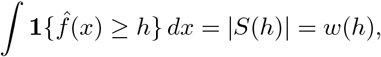

we obtain

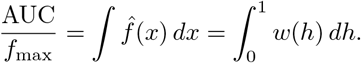

Therefore, AUC/*f*max is not the width at a single threshold. Instead, it is the integral of the profile width over all normalized intensity thresholds. This explains why AUC/*f*max can capture broader shape information than FWHM, which only measures the width at half-maximum.

For an ideal Gaussian intensity profile,

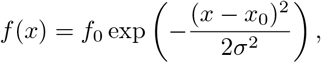

the maximum intensity is

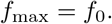

The area under the curve is

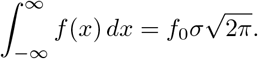

Thus,

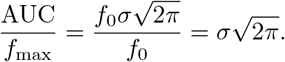

For the same Gaussian profile, the full-width-at-half-maximum is

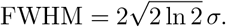

Solving for *σ* and substituting into the AUC/*f*max expression gives

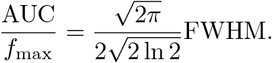

Therefore,

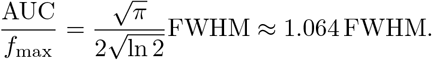

Thus, for a perfect Gaussian profile, AUC/*f*max is directly proportional to FWHM and yields a similar length measurement. However, AUC/*f*max remains sensitive to the full normalized 1D kinetochore intensity profile, without requiring it to be Gaussian, making it a more versatile measure of kinetochore length.

